# *TheWeight*: A simple and flexible algorithm for simulating non-ideal, age-structured populations

**DOI:** 10.1101/2022.01.05.475135

**Authors:** Robin S. Waples

## Abstract

1. The Wright-Fisher model, which directs how matings occur and how genes are transmitted across generations, has long been a lynchpin of evolutionary biology. This model is elegantly simple, analytically tractable, and easy to implement, but it has one serious limitation: essentially no real species satisfies its many assumptions. With growing awareness of the importance of jointly considering both ecology and evolution in eco-evolutionary models, this limitation has become more apparent, causing many researchers to search for more realistic simulation models.
2. A recently described variation retains most of the Wright-Fisher simplicity but provides greater flexibility to accommodate departures from model assumptions. This generalized Wright-Fisher model relaxes the assumption that all individuals have identical expected reproductive success by introducing a vector of parental weights ***w*** that specifies relative probabilities different individuals have of producing offspring. With parental weights specified this way, expectations of key demographic parameters are simple functions of ***w***. This allows researchers to quantitatively predict the consequences of non-Wright-Fisher features incorporated into their models.
3. An important limitation of the Wright-Fisher model is that it assumes discrete generations, whereas most real species are age-structured. Here I show how an algorithm (*TheWeight*) that implements the generalized Wright-Fisher model can be used to model evolution in age-structured populations with overlapping generations. Worked examples illustrate simulation of seasonal and lifetime reproductive success and show how the user can pick vectors of weights expected to produce a desired level of reproductive skew or a desired *N_e_/N* ratio. Alternatively, weights can be associated with heritable traits to provide a simple, quantitative way to model natural selection. Using *TheWeight*, it is easy to generate positive or negative correlations of individual reproductive success over time, thus allowing explicit modeling of common biological processes like skip breeding and persistent individual differences.
4. Code is provided to implement basic features of *TheWeight* and applications described here. However, required coding changes to the Wright-Fisher model are modest, so the real value of the new algorithm is to encourage users to adopt its features into their own or others’ models.

## 1 INTRODUCTION

The Wright-Fisher (WF) model of reproduction is a lynchpin of evolutionary biology, as it describes how matings occur and how genes are transmitted across generations. This model also provides a direct link between the ecological consequences of abundance, which depend on census size (*N*), and the evolutionary consequences of abundance, which depend on effective population size (*N_e_*). When populations dynamics follow the standard WF model, the number of offspring per parent is approximately Poisson distributed (with variance ≈ mean), in which case the census and effective sizes are the same. Since the time of the Modern Synthesis (Huxley 1942), theoretical population genetics has played a seminal role in shaping the development of evolutionary biology. Virtually every equation in population genetics includes a term for effective size, and it is routine to adopt the assumptions of the WF model to take advantage of this equivalence between *N* and *N_e_.* Under WF assumptions (a single closed population with discrete generations, constant size, random mating and random variation in reproductive success), the loss of genetic diversity due to drift depends only on the number of reproductive individuals, so both demographic and evolutionary processes can be modeled with a single parameter for population size. A population that satisfies all of these assumptions is said to be “ideal.” These features make the WF model easy to simulate in a computer, and as a consequence it is incorporated into a wide variety of software for forward-in-time and coalescent simulations (Peng et al. 2015; NCI 2018).

Despite its advantages and nearly universal appeal, the standard WF model has an important limitation: its many assumptions are compatible with essentially zero species in nature. Almost all real populations deviate from WF assumptions in ways that cause the effective size to be less than the number of adults, primarily because of skewed sex ratio and greater-than-Poisson variance in offspring number within sexes, and these deviations lead to a wide range of *N_e_/N* ratios in nature (Frankham 1995; Palstra and Fraser 2012). From the evolutionary perspective this might not be a problem, as a researcher who wants to model genetic processes in a real population with census and effective sizes of, say, *N* = 1000 and *N_e_* = 200, could just model a WF population with *N* = *N_e_* = 200. However, populations with *N* = 200 and *N* = 1000 generally are not equivalent for ecological processes such as competition, predation, and individual behavior, so exclusive reliance on WF assumptions constrains the ability of a model to adequately capture realistic eco-evolutionary dynamics. In recent decades it has become apparent that most problems in biology are not strictly ecological or evolutionary but rather are eco-evolutionary in nature (Pelletier et al. 2009), and researchers have increasingly expressed the desire for more realistic models for population genetic simulations (Haller and Messer 2019).

Because the WF model requires numerous simplifying assumptions, there are many ways to model non-ideal populations (e.g., Eldon and Wakeley 2006; Der et al. 2011; Chotibut and Nelson 2017). A common approach to incorporate overdispersed variation in reproductive success is to draw offspring numbers from a negative binomial or gamma distribution, which can be scaled to produce a desired level of reproductive skew. This approach, however, is better suited to modeling population-level processes than characteristics of individuals. Here I describe in detail an algorithm, *TheWeight*, which makes a simple change to the WF model that provides considerable added flexibility to simulate realistic, individual-based demographics. In the standard WF model, every individual has an equal chance to produce each offspring; in *TheWeight*, the user defines a vector of parental weights that specifies the relative probabilities that each individual will be chosen as a parent for a given offspring. Individual weighting schemes have been incorporated into a number of genetic simulators, such as *Nemo* (Guillaume and Rougemont 2006), *simuPOP* (Peng and Amos 2008), and *SFS_CODE* (Hernandez 2008). *TheWeight* takes advantage of recent theoretical developments that show that many key demographic parameters can be predicted based on simple functions of the vector of parental weights. However, these results, and the original description of the generalized Wright-Fisher model (Waples 2020), followed the traditional WF model in assuming that generations are discrete.

Most real populations are age structured, so the discrete-generation assumption is an important limitation. Here I illustrate how *TheWeight* algorithm can be used to model eco-evolutionary processes in iteroparous species with overlapping generations. The treatment here focuses on the large fraction of species that exhibit strongly seasonal, birth-pulse reproduction (Caswell 2001). For these species, reproductive success is commonly assessed from two different perspectives: seasonal (in which offspring number is the total produced by an individual across a reproductive season—here assumed to be years) and lifetime (in which lifetime reproductive success (*LRS*) is the cumulative number of offspring produced across an individual’s lifespan). Variance in *LRS* has received considerable attention (Hill 1972; Brown 1988; Kruuk et al. 1999; Tuljapurkar et al. 2020; Snyder et al. 2021) because it integrates information across generational time scales and is the most important factor determining effective population size per generation (*N_e_*). However, analysis of data on seasonal reproduction provides important insights into mating systems. Furthermore, seasonal reproduction is much easier to study, especially in long-lived species, and many monitoring programs routinely estimate the effective number of breeders per year (*N_b_*) and other population genetic parameters using seasonal data (Pudovkin et al. 1996; Ruzzante et al. 2016; Ackerman et al. 2017).

Worked examples illustrate how *TheWeight* can be used to model (1) changes in fecundity with age; (2) variation in expected reproductive success among individuals of the same age and sex; and (3) positive or negative correlations in individual reproductive success over time. This latter feature is useful for modeling common biological features such as intermittent breeding and persistent individual differences (aka “individual quality”; Wilson and Nussey 2010). Using *TheWeight*, it is also easy to simulate null models for age-structured species that incorporate random survival and/or random variation in reproductive success; these null models in turn provide important context for evaluating empirical data. Computer code is provided to illustrate how these features can be implemented, but the required coding changes are simple enough that users can easily incorporate them into their own or others’ models.

## 2 METHODS

### 2.1 Wright-Fisher reproduction

#### 2.1.1 The standard Wright-Fisher model

The standard Wright-Fisher model has the following features: population size is constant; generations are discrete; selection is absent; mating is random; and every individual is equally likely to be the parent of a given offspring, regardless how many or few offspring that individual has already produced. Conceptually, reproduction occurs as follows: each individual contributes an equal and very large number of gametes to a huge (essentially infinite) pool of gametes, which unite at random to produce zygotes. Random survival of zygotes then produces the few individuals that form the next generation (Wright 1931). A population meeting all of these criteria is said to be “ideal” and has the property that the effective population size (*N_e_*) equals (on average) the adult census size (*N*).

This latter feature can be illustrated using standard definitions of inbreeding *N_e_* based on the mean (*μ_k_*) and variance 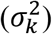 among parents in number of offspring (*k*) (Crow and Denniston 1988):

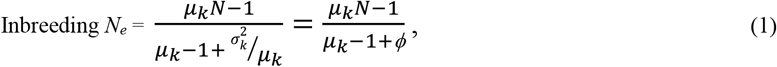

where 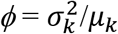 is the ratio of variance to mean offspring number. Focus is on inbreeding *N_e_* because it relates to the number of parents and hence is more relevant than variance *N_e_* to the topics considered here. Furthermore, inbreeding *N_e_* is much less sensitive to effects of sampling and hence is more flexible than variance *N_e_*, especially for applications involving age-structured populations (Waples 2002, 2020). Equation 1 applies to monoecious/hermaphroditic diploids with random selfing; slight variations are available for other mating systems. Implementing the WF model leads to multinomial sampling of parents, so 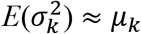. Inserting *ϕ* = 1 into Equation 1 yields the result that *N_e_* ≈ (*μ_k_N* – 1)/*μ_k_* ≈ *N*. [The result is exact if one uses the exact (binomial) variance 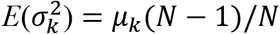 rather than the Poisson approximation; see Supporting Information.]

The WF model is easy to implement in a computer, the only required input parameters being the number of ideal individuals (*N*) and, if genotypes are modeled, the number of gene loci being tracked. Then, only three more steps are required to complete one generation of reproduction (Figure 1A). In Step 2, each offspring “chooses” its parents randomly and with replacement from the *N* potential parents. In Step 3, at each gene locus one of the two alleles from each parent is randomly chosen to pass on to the offspring. Step 4 completes the cycle, as offspring become potential parents of the next generation.

**Figure 1.**
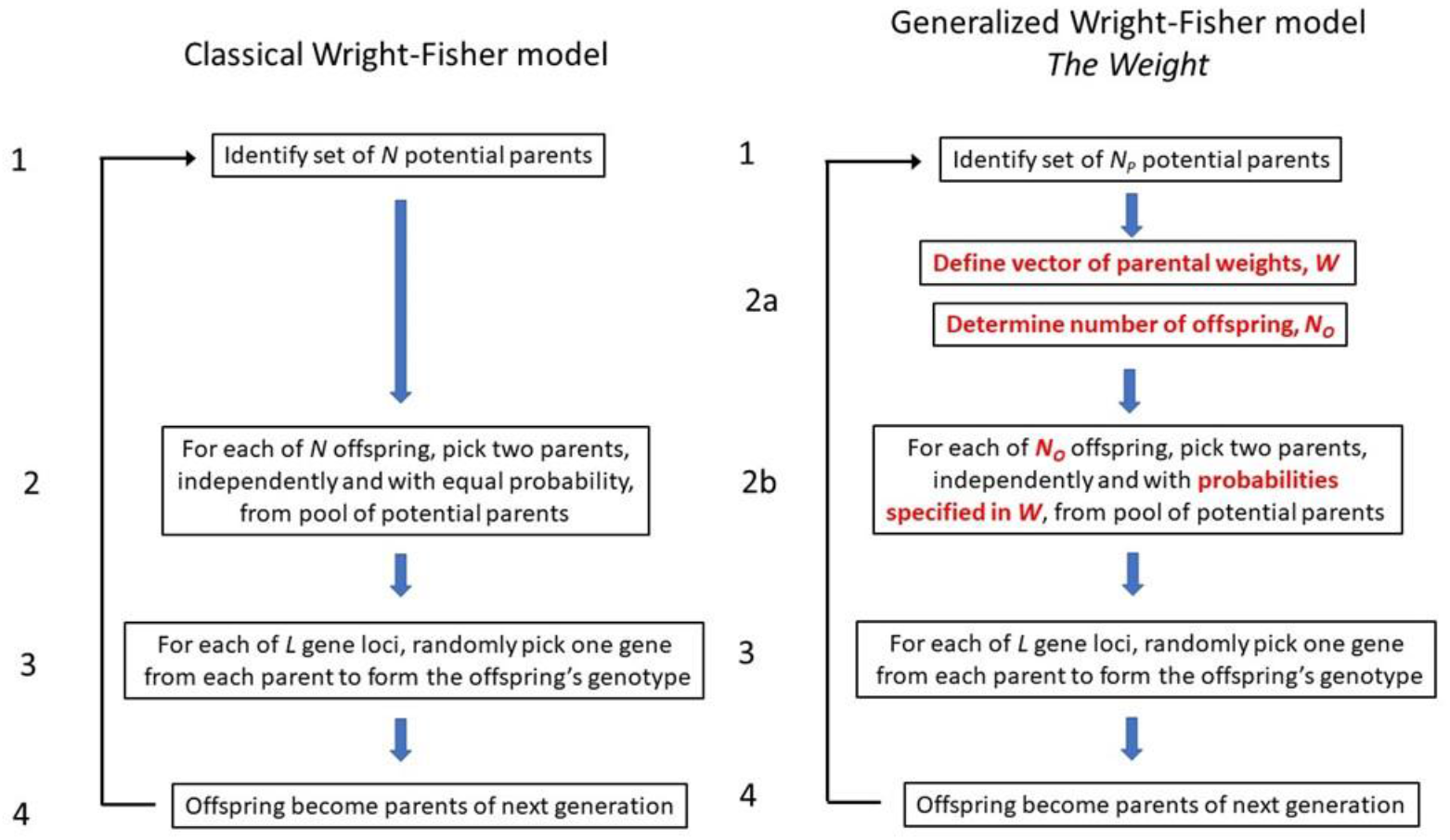
Flowcharts showing algorithms for the WF model (left) and *TheWeight* (right), with differences highlighted in **bold red**.

#### 2.1.2 The generalized Wright-Fisher model

Conceptually, the generalized WF model differs in one important way from the standard version: the parents are allowed to make unequal contributions to the initial gene pool, with the contributions being proportional to the weight (*wi*) for each parent. The first use of a weighted WF model like this was by Robertson (1961) for the special case of full-sibling families, and the model was further developed by Waples (2020). Let *μ* and 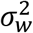 be the mean and variance, respectively, of the vector of parental weights, and let *CV_w_* = *σ_w_*/*μ_w_* be the coefficient of variation of ***w***. [*CV_w_* is the same when calculated using the raw parental weights ***w*** or weights that are standardized (e.g., so that they sum to 1 or have mean = 1), so here the raw weights are used]. Then, several key population genetic parameters can be expressed as simple functions of the squared *CV* of the parental weights, 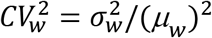:

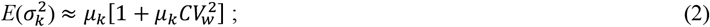

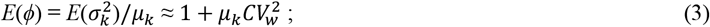

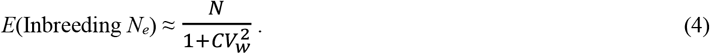

These are Equations 12, 13, and 20, respectively, from Waples (2020). Equations 2 and 3 are conditional expectations that depend on mean offspring number; they are expressed in terms of population parameters 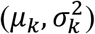 but also are unbiased as conditional expectations for samples (Waples 2020). Equation 4 is a general result for inbreeding *N_e_* that does not depend on mean offspring number (Waples 2020); for variance *N_e_*, this equation holds only if *μ_k_* = 2 (Felsenstein 2019). Equation 4 can be converted into an expression that predicts the *N_e_/N* ratio. If one divides both sides by *N*, the result is:

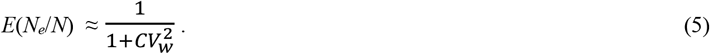

When all parental weights are equal, *CV_w_* = 0 and Equation 5 predicts *N_e_/N* = 1, as expected for the standard Wright Fisher model.

In computer modeling, the only difference between the traditional and generalized WF models occurs at Step 2: instead of choosing adults with equal probability to be parents of a given offspring, the probability a parent will be chosen is proportional to its weight (Figure 1B). To deal effectively with age structure, it is also important to allow the overall number of offspring to vary from the constraint in the WF model that *μ_k_* = 2. Modeling separate sexes requires separate routines for choosing male and female parents, together with separate vectors of parental weights. Effective size can then be calculated separately for males and females (*N_em_, N_ef_*), and overall *N_e_* can be obtained using Wright’s (1938) sex ratio adjustment: *N_e_* = 4*N_em_N_ef_* /(*N_em_* + *N_ef_*).

### 2.2 Modeling age-structured populations with *TheWeight* algorithm

#### 2.2.1 Seasonal reproduction

##### 2.2.1.1 Age-specific vital rates

For seasonal reproduction in a discrete-time, birth pulse model, age structure can be modeled using age-specific vital rates from a life table: *s_x_* = probability of surviving from age *x* to *x*+1, *b_x_* = mean number of offspring produced by an individual of age *x*, and *N_x_* = number of individuals in age class *x*. In an age-structured model based on *TheWeight*, a simple option is to assign parental weights based on the *b_x_* value for the appropriate age. If all individuals of the same age receive the same weight, then each age class behaves like a mini WF population, with the expected age-specific variance in offspring number being approximately equal to the mean: 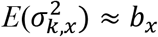. In that case, the expected value of the variance-to-mean ratio (*ϕ_x_*) is 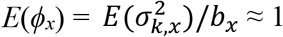.

Assuming that *ϕ_x_* = 1 for each age and sex might be reasonable for some populations, but that is not universally the case. Empirical data for age-specific *ϕ_x_* are seldom published, but (for example) for black bears in Michigan, USA, *ϕ_x_* was significantly >1 for both females and males and estimated to be 10 or higher for males aged 6-10 years (Waples et al. 2018). A more comprehensive life table would then include another age-specific vector of vital rates: *ϕ_x_*.

To model the desired level of overdispersion at each age (indicated by *ϕ_x_*), the following steps can be used to generate *N_x_* parental weights. This process can be done separately for male and female parents of each offspring.

Step 1. Determine the target value for the squared *CV* of parental weights. By rearranging Equation 3,

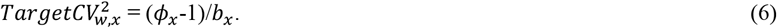
Step 2. For any given 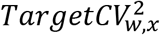, it is possible to identify a Poisson distribution with mean = variance = *λ_x_* that will produce a vector of parental weights such that 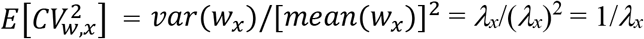. The required mean of the Poisson distribution is thus 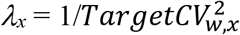.
Step 3. Simulating *N_x_* random Poisson values using parameter *λ_x_* will thus, on average, produce a vector of parental weights with the desired 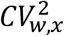. This process can be iterated until the realized 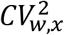 is within a specified tolerance of the target (most results presented here used tolerance = 1%).

With the parental weights specified for each age, seasonal reproduction in the generalized WF model can proceed using either a two-step or one-step process.

Step 4A. In the two-step process, for each offspring one first determines the age of the parent(s). This can be done randomly (e.g., by sampling the parent’s age from the ages of the adult lifespan, with relative probabilities being the product *N_x_b_x_*). Next, the actual parent can be chosen randomly using the appropriate age-specific vector of parental weights generated in Step 3.
Step 4B. In the single-step process, adults of all ages (and their weights) are combined into a single matrix, and the parent(s) for each offspring are chosen based on the combined vector of parental weights. This requires that the mean weight for each age 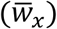 be proportional to the age-specific mean fecundity (*b_x_*). If *b_x_* is constant, the weighting scheme described above already accomplishes this. More generally, if *b_x_* varies with age, then for each age every parental weight has to be rescaled by multiplying by the factor *F* = *b_x_/λ_x_.* After rescaling, 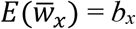 for each age; although both the mean and variance of the weights have changed, the squared *CV* remains the same.

##### 2.2.1.2 Natural selection

Stabilizing selection is commonly modeled using a Gaussian fitness function applied to a continuous phenotypic trait *z,* such as body size (Lande 1976). A general way to implement this using *TheWeight* is as follows:

Step 1: Define the phenotypic distribution, *p*(*z*). *z* is commonly assumed to have a normal distribution with mean = *μ_z_* and 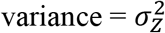. In that case,

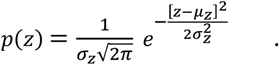
Step 2: Map the phenotypes to the fitness function to produce the distribution of relative individual fitness *p*(*w*), which is identical to the distribution of parental weights. Under stabilizing selection, the maximum fitness (*w_max_*) occurs at an intermediate phenotype, denoted by *θ,* such that individual fitness values (*w_i_*) are a function of the squared distance between the individual’s phenotype and *θ*.

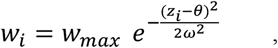

where *ω* = *σ_w_* is the “width” of the fitness surface.

Steps 1 and 2 are commonly used to model selection. The advantage of using *TheWeight* is that the expected consequences of selection for key demographic and population genetic parameters can be quantified.

Equation 3 above shows that the expected value of the variance to mean ratio for offspring number 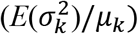 is a simple function of the squared *CV* of parental weights. A related quantity defined by Crow (1958) is 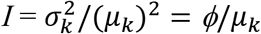. Commonly referred to as the Opportunity for Selection, *I* is the variance in relative fitness, which places an upper limit on the evolutionary response to selection (Walsh and Lynch 2018). Dividing each side of Equation 3 by the mean offspring number produces this result:

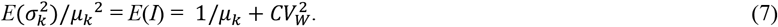

If the phenotypic distribution is continuous, the distributions of fitnesses and parental weights are also continuous, in which case the means and variances can be calculated using the following formulas (lower bounds of the integrals are set to 0 because fitnesses and parental weights cannot be negative):

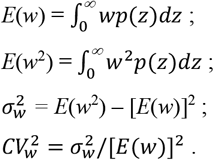

The above example assumed a Gaussian distribution of selection coefficients, but this approach can also be used more generally with any selection function (linear, quadratic, etc.). Equations 2-5 and 7 do not depend on the full distribution of parental weights—only the *CV*.

##### 2.2.1.3 Generic weighting schemes

Generic weighting schemes that assign weights randomly to individuals are easy to implement. One way to do this is to randomly assign potential parents ordered cardinal numbers *i* = 1, 2,… *N* and then give each a relative weight that is a function of their assigned number. Simple examples include power functions (*w*_i_ = *i^x^* = 1^*x*^, 2^*x*^, … *N^x^*) and harmonic functions (*w*_i_ = 1/*i^x^* = 1/1^*x*^, 1/2^*x*^, … 1/*N^x^*). Equations 2-5 can be used to predict the consequences of these weighting schemes for parameters of interest.

#### 2.2.2 Lifetime reproductive success

Analysis of *LRS* focuses on a birth cohort of individuals that are enumerated at a specified age (here assumed to be age at maturity). *TheWeight* has three general options for modeling lifetime reproductive success. The simplest approach is to generate new weights for each age to match the species’ vital rates (as described in Section 2.2.1.1) and then assign those randomly to the survivors each year. This approach is suitable if it is reasonable to assume that an individual’s reproductive success is uncorrelated over time (as assumed by Felsenstein 1971, Hill 1972, and Waples et al. 2011). *TheWeight* can also be used to model two common scenarios in which this independence assumption does not hold.

##### 2.2.2.1 Intermittent breeding

In many species, females (and sometimes males) do not necessarily breed every year, leading to a strategy that has been called “skip breeding” (Goutte et al. 2011) or “intermittent breeding” (Shaw and Levin 2011). This type of life history can be found in vertebrate ectotherms, mammals, birds, and some invertebrates; it can lead to positive or negative correlations in the number of offspring an individual produces over time, depending on the time lag and pattern of skip breeding. This life history can be described by a vector *θ,* the *t*^th^ element of which (*θ_t_*) is the probability that an individual will reproduce in the current year, given that it last reproduced *t* years before (Shaw and Levin 2011). For example, Waples and Antao (2014) estimated that *θ_t_* = [0.025, 0.443, 0.634, 0.743, 1] for female loggerhead turtles, reflecting substantial costs of migration and reproduction. In contrast, for female black bears in Michigan *θ_t_* = [0.035, 0.94, 1] (Waples et al. 2018), indicating a faster recovery of reproductive potential. This life-history can easily be modeled with *TheWeight*.

Step 1: For each individual, determine the value of *t* = the number of years since the individual last reproduced.
Step 2: With probability *θ_t_,* replace that individual’s weight for the current year with 0.

##### 2.2.2.2 Persistent individual differences

Persistent individual differences in reproductive success occur when some individuals are consistently above or below average (for their age and sex) at producing offspring. In iteroparous species, persistent individual differences increase lifetime variance in offspring number and reduce *N_e_* (Lee et al. 2011, 2020). Variation in size at age can easily lead to this type of pattern: an individual that is larger (and hence more fecund) than others in its cohort at age *x* is also likely to have above-average size and fecundity at older ages. Persistent differences could have a genetic basis or might simply reflect early-life-history environmental “silver spoon” effects with long-term consequences.

This life history feature can be modeled in *TheWeight* by introducing positive correlations between parental weights over time, which can be accomplished by retaining individual weights across lifetimes. However, because the target value of desired 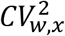 is contingent on *b_x_* and *ϕ_x_*, when these vital rates change with age the squared *CV* of the parental weights must also change. Doing this without distorting the relative rankings of the individual weights is tricky but can be accomplished as follows:

Step 1. Using the approach outlined in the previous section, generate a vector of parental weights ***W***_1_ for the *N_1_* age-1 individuals in a cohort, for which expected fecundity is *b_1_* and 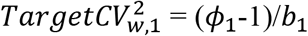. The Poisson parameter to produce the desired parental weights at age 1 is 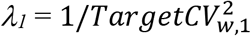. If age-1 individuals are not mature, start this process with the age at sexual maturity.
Step 2. Now we need a vector of weights ***w***_2_ that retain the same framework of individual differences but produce 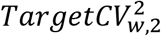 at age 2 = (*ϕ*_2_-1)/*b*_2_. This can be achieved by adding a constant *C*_2_ to each individual weight generated at age 1. This has no effect on the variance of the weights but changes the mean, so it also changes the squared *CV*.
Step 3. Define the factor 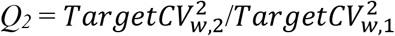. Then, 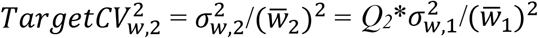. Now 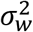 is the same at ages 1 and 2, so this last equation implies that 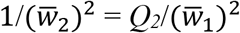 and hence 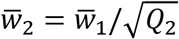.
Step 4. If the increase in the mean weight is accomplished by adding a constant, it is also true that 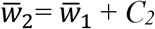. Solving these two equation yields the desired result:

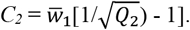
Step 5. Repeat this process for each additional year of the maximum adult lifespan (*AL* years). The result is a *N_1_* x *AL* matrix of parental weights. Each year, weights for the survivors in a cohort are taken from the appropriate column of the matrix.

Adding a constant to all individual weights changes their relative values. However, it maintains the same rank order of weights, and the pairwise correlations of the age-specific vectors of individual weights are all *r* = 1.0. Weaker positive correlations could be introduced by adding a random normal error term to the individual weights each year without shuffling them.

### 2.3 Simulations

To illustrate calculations described above and evaluate their accuracy, seasonal and lifetime reproductive success was simulated using *TheWeight* algorithm. In addition, one scenario simulated multilocus geneotypes and tracked loss of heterozygosity over time. Simulations were conducted in R (R Core Team 2021) using code available on Zenodo.

## 3 RESULTS

### 3.1 Seasonal reproduction

#### 3.1.1 Modeling age-specific vital rates

Table 1 illustrates how to compute the vectors of parental weights when modeling seasonal reproduction according to age-specific vital rates from an expanded life table, which also includes age-specific values for *ϕ* for a single sex, nominally male. The hypothetical species is first enumerated at age 1, matures at age 3, and has a maximum lifespan of 10 years. With annual survival = 0.7 and a fixed cohort size of 1000 age-1 males, expected numbers of males alive at subsequent age are given by the vector *N_x_*. Expected fecundity (*b_x_*) is proportional to age and is scaled to production of a stable population (which produces 2000 yearling offspring per year, 1000 of each sex). In this example, the overdispersion index *ϕ* was also chosen to increase with age, but more slowly than fecundity. For each age, Table 1 shows how to calculate the 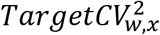 that will produce the desired level of overdispersion, and the respective Poisson parameter that can be used to simulate random weights with the desired properties. When using the two-step process to simulate reproduction (Step 4A in Section 2.2.1.3), the age(s) of the parent(s) of each offspring are first chosen, and then the actual parent is chosen from that age group, with sampling probabilities determined by the parental weights. With this two-step approach, the initial Poisson parameters (*Λ_x_*) can be used for each age. In this example, however, reproduction is modeled in a single step (as in Step 4B in Section 2.2.1.3), and the weights for each age have to be rescaled so that age-specific 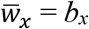. This is accomplished by multiplying the initial *Λ_x_* by an age-specific scaling factor. Across 1000 replicate simulated populations, mean realized *ϕ_x_* for each age closely agreed with the parametric expectations (Table 1).

**Table 1.** A life table for one sex (nominally male) of a hypothetical species, illustrating how to generate the desired level of reproductive skew in seasonal reproduction, using parental weights. Each cohort begins with 1000 age-1 males; annual survival of 0.7 and maturity at age 3 produces a total of *N_A_* = 1539 adult (age 3+) males. For each age *x*, population parameters are *N_x_* = fixed number of males alive; *b_x_* = expected fecundity; *ϕ_x_* = expected ratio 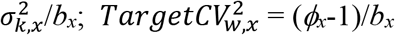; initial 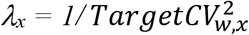; scaling factor = *b_x_*/(initial *Λ_x_*); rescaled *Λ_x_* = (initial *Λ_x_*)*scaling factor = *b_x_*. Observed values of *ϕ_x_*, 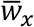, and 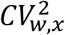 are averages across 1000 replicate simulations.

| Age | $N_x$ | $b_x$ | $\phi_x$ | Target<br>$CV_{w,x}^2$ | initial<br>$\lambda_x$ | scaling<br>factor | rescaled<br>$\lambda_x$ | Observed | | |
| --- | --- | --- | --- | --- | --- | --- | --- | --- | --- | --- |
| | | | | | | | | $\phi_x$ | $\bar{w}_x$ | $CV_{w,x}^2$ |
| 1 | 1000 | 0 |  |  |  |  |  |  |  |  |
| 2 | 700 | 0 |  |  |  |  |  |  |  |  |
| 3 | 490 | 0.805 | 3.00 | 2.484 | 0.403 | 2.000 | 0.805 | 2.99 | 0.805 | 2.483 |
| 4 | 343 | 1.073 | 3.25 | 2.097 | 0.477 | 2.250 | 1.073 | 3.23 | 1.073 | 2.099 |
| 5 | 240 | 1.342 | 3.50 | 1.863 | 0.537 | 2.500 | 1.342 | 3.49 | 1.342 | 1.865 |
| 6 | 168 | 1.610 | 3.75 | 1.708 | 0.585 | 2.750 | 1.610 | 3.73 | 1.610 | 1.703 |
| 7 | 118 | 1.878 | 4.00 | 1.597 | 0.626 | 3.000 | 1.878 | 3.94 | 1.878 | 1.598 |
| 8 | 82 | 2.146 | 4.25 | 1.514 | 0.660 | 3.250 | 2.146 | 4.26 | 2.146 | 1.515 |
| 9 | 58 | 2.415 | 4.50 | 1.449 | 0.690 | 3.500 | 2.415 | 4.54 | 2.415 | 1.451 |
| 10 | 40 | 2.683 | 4.75 | 1.398 | 0.715 | 3.750 | 2.683 | 4.78 | 2.683 | 1.397 |
| ----- |  |  |  |  |  |  |  |  |  |  |
| Total $N$ | 3239 | | | | | | | | | |
| Adult $N$ | 1539 | | | | | | | | | |

Most life tables do not contain data for the age-specific parameter ϕ_x_, so it is common to assume that all ϕ_x_ = 1. The consequences of violation of this assumption can easily be modeled using *TheWeight* by replacing empirical ϕ_x_ estimates with 1.0. This was done for the vital rates in Table 1, with results shown in Table S1. Across parents of all ages, assuming all ϕ_x_ = 1 had no effect on the overall mean offspring number (*μ_k_* = 1.30), but it reduced overall 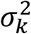 from 5.00 to 1.56 and overall ϕ from 3.85 to 1.20—both reductions of almost 70%.

These examples used data for the entire population. Subsampling offspring is easy to accomplish using *TheWeight*, as illustrated in the R code provided.

#### 3.1.2 Modeling selection

In modeling seasonal reproduction using the vital rates shown in Table 1, given that across all adults *μ_k_* = 1.30 and mean 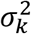was 5.00 (Table S1), the overall annual Opportunity for Selection was 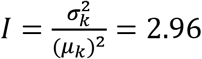. Interpretation of raw values of *I* is complicated by its dependence on mean offspring number, which can reflect experimental design and sampling effort more than underlying biology (Downhower et al. 1987; Fairbairn and Wilby 2001). A simple way to resolve this problem is to subtract the expected contribution from random demographic stochasticity (1/*μ_k_*), which produces an adjusted Opportunity for Selection that is independent of mean fitness (Waples 2020):

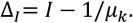

From Equation 7, 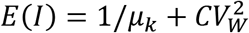, so the expected value of *Δ_I_* is simply the squared coefficient of variation of the parental weights:

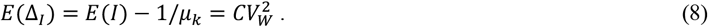

The adjusted Opportunity for Selection index, which can be calculated directly from the distribution of relative fitness, represents the component of variance in relative fitness that exceeds that expected under a null model of random reproductive success.

For the vital rates shown in Table 1, mean *I* = 2.96 so Δ_*I*_ = 2.96 – 1/1.30 = 2.19, which was also the mean value of the actual 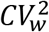 in the simulations (Table S1). Because Δ_*I*_ does not depend on mean offspring number, it can be directly compared to similar values obtained in other studies. Knowing 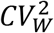 also allows one to predict the expected reduction in the annual *N_b_/N* ratio that can be attributed to natural selection. For this example, based on Equation 5, 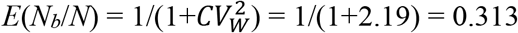, indicating that unequal expectations for reproductive success attributed to selection would be expected to drop the effective number of breeders to less than one third of the number of adults. In contrast, if all ϕ_x_ were assumed to be 1 (as in Table S1), 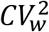 would shrink dramatically (to 0.15), with the remaining variation in parental weights being reduced to only the among-age effect (variation in *b_x_* with age). Under that scenario, the expected value of *N_b_/N* would be 1/(1.15) = 0.87, for a modest reduction of 13%.

It should be noted that although in some cases parental weights can be equated directly to selection coefficients (as described in Section 2.2.1.2), individuals can have unequal expectations of reproductive success for a variety of reasons that are more related to luck than to pluck (Snyder and Ellner 2018) or individual quality (Wilson and Nussey 2010). As shown in Equation 7, variation in parental weights is more directly related to the concept of the Opportunity for Selection, which might not be a reliable indication of the actual strength of selection (Downhower et al. 1987; Fairbairn and Wilby 2001; Jennions et al. 2012).

#### 3.1.3 Generic weighting schemes

The consequences for the *N_e_/N* ratio of applying some simple rules for specifying parental weights are shown in Figure 2. In the first three scenarios, the weights were power functions that produced increasing amounts of reproductive skew: ***w*_i_** = *i*^1^, *i*^2^, and *i*^3^. The next three scenarios used the analogous harmonic series: ***w*_i_** = 1/*i*^1^, 1/*i*^2^, and 1/*i*^3^; and the final scenario used an exponential series: ***w*_i_** = *e^i^*. It is apparent from Figure 2 that the power functions produce *N_e_*/*N* ratios that are largely insensitive to *N*, whereas for the other series log(*N_e_*/*N*) declines approximately linearly with log(*N*). The number of possible weighting schemes is effectively unlimited, but these simple examples show that it is easy for users to pick a weighting scheme that will produce essentially any desired *N_e_/N* ratio or any desired Opportunity for Selection.

**Figure 2.**
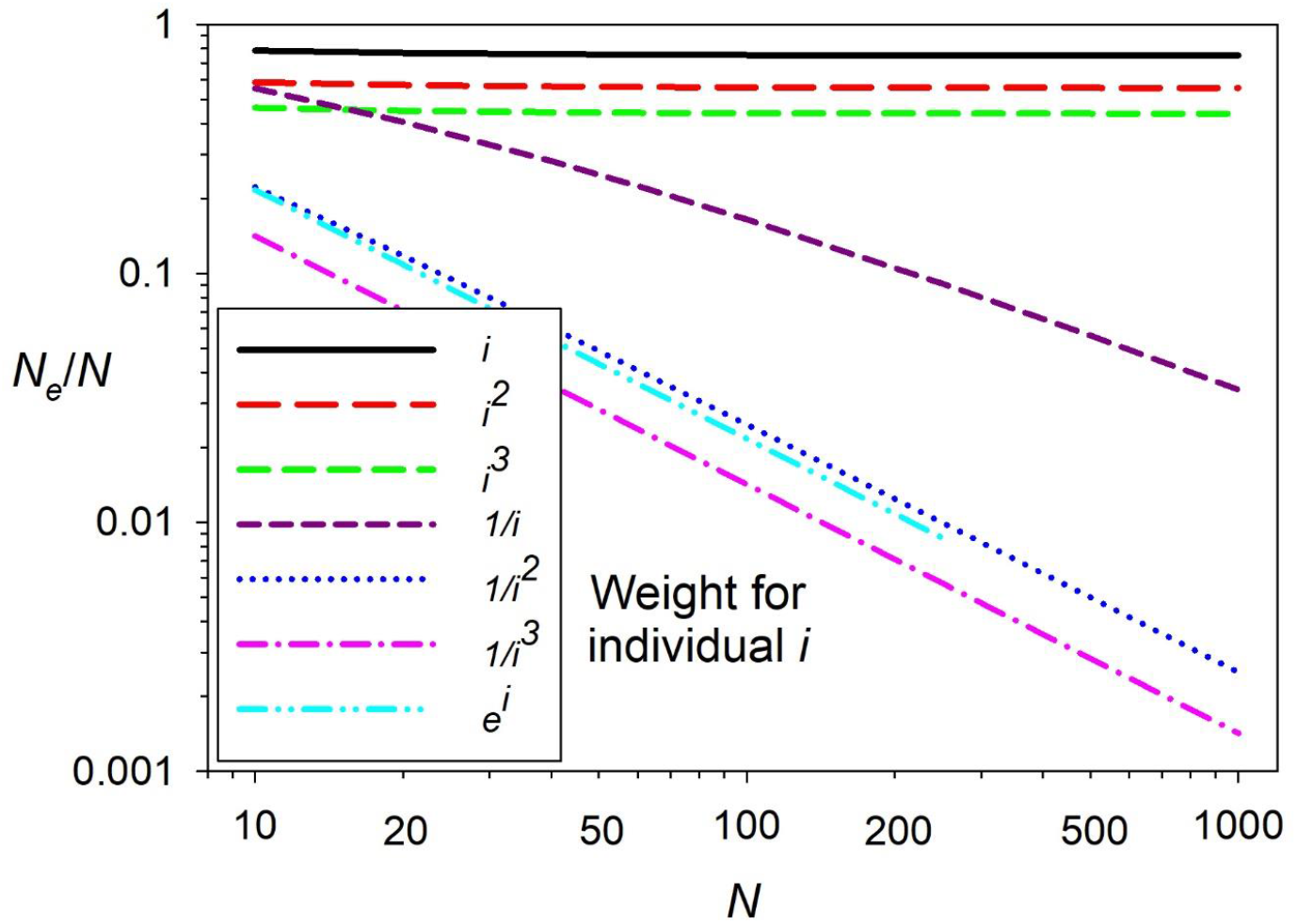
Predicted effects of seven different parental weighting schemes on the ratio *N_e_/N* based on Equation 5. For each individual *i*, *i*=1,*N*, the legend shows the formula for computing the relative parental weight ***w*_i_**. Overflow issues in R prevent showing results for *e^i^* when *i* > 250.

### 3.2 Lifetime reproductive success

Table 2 uses the vital rates in Table 1 to illustrate how to generate parental weights that produce persistent individual differences in reproductive success over time. Since ages 1 and 2 are juveniles (and hence have zero weights), this table begins with age at maturity = 3 years. First, the 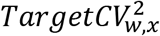 expected to produce weights with the desired level of overdispersion at each age is computed as (*ϕ_x_*-1)/*b_x_*, and the Poisson parameter to generate weights for age-3 adults is then 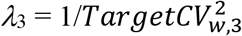—all as in Table 1. Next, age-specific constants are calculated as 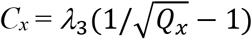, where 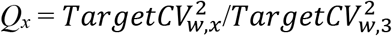, and adjusted parental weights for subsequent ages are calculated as 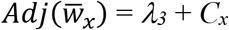. Results of the simulations show that although these adjustments changed both the means and variances of the age-specific parental weights, they did not change 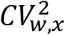, and the actual mean *ϕ_x_* values were very close to the parametric expectations for each age (Table 2), just as they were for the example in Table 1. The adjusted weights, however maintain the relative weights for each individual over time.

**Table 2.** Example illustrating how to generate parental weights to model persistent individual differences in lifetime reproductive success, for a cohort of 490 males that survive to age at maturity (3). The first six columns repeat adult data from Table 1. The next three columns show calculation of adjustments to individual age-3 weights to generate weights for subsequent ages. These adjusted weights produce the target 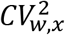 for each age while maintaining the age-3 pattern of relative individual fitness. *λ*_3_ is the Poisson parameter that will generate weights with the desired properties for age-3 adults; 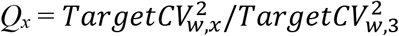; 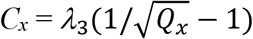; 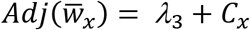; Observed *ϕ_x_* = means across 1000 replicate simulations.

| Age | $N_x$ | $b_x$ | $\phi_x$ | Target<br>$CV_{w,x}^2$ | $\lambda_3$ | $Q_x$ | $C_x$ | $Adj(\bar{w}_x)$ | Observed<br>$\phi_x$ |
| --- | --- | --- | --- | --- | --- | --- | --- | --- | --- |
| 3 | 490 | 0.805 | 3.00 | 2.484 | 0.403 | 1.000 | 0.000 | 0.403 | 3.00 |
| 4 | 343 | 1.073 | 3.25 | 2.097 | - | 0.844 | 0.036 | 0.438 | 3.26 |
| 5 | 240 | 1.342 | 3.50 | 1.863 | - | 0.750 | 0.062 | 0.465 | 3.50 |
| 6 | 168 | 1.610 | 3.75 | 1.708 | - | 0.688 | 0.083 | 0.485 | 3.76 |
| 7 | 118 | 1.878 | 4.00 | 1.597 | - | 0.643 | 0.099 | 0.502 | 4.03 |
| 8 | 82 | 2.146 | 4.25 | 1.514 | - | 0.610 | 0.113 | 0.516 | 4.25 |
| 9 | 58 | 2.415 | 4.50 | 1.449 | - | 0.583 | 0.124 | 0.527 | 4.49 |
| 10 | 40 | 2.683 | 4.75 | 1.398 | - | 0.563 | 0.134 | 0.537 | 4.79 |

Once the matrix of lifetime parental weights is established, it requires only a single line of code to switch between modeling reproductive success that is independent and perfectly correlated over time: for independence the parental weights are randomly scrambled every year, and to model persistent individual differences they are not. This can make a large difference in terms of reproductive skew: with parental weights randomized each year, the median variance in *LRS* across 1000 simulations for the 490 males in each cohort that reached age 3 was 30.7; with parental weights retained across lifetimes, median 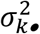 more than doubled, to 76.1. Persistent individual differences also greatly skewed the distribution of *LRS* compared to a null model of independence (Figure 3). In simulations for which parental weights were retained, almost half of the males that reached age at maturity never produced an offspring, while some others produced 100 or more. When weights were shuffled each year, maximum *LRS* was <60 and null parents comprised a bit over a third of each cohort.

**Figure 3.**
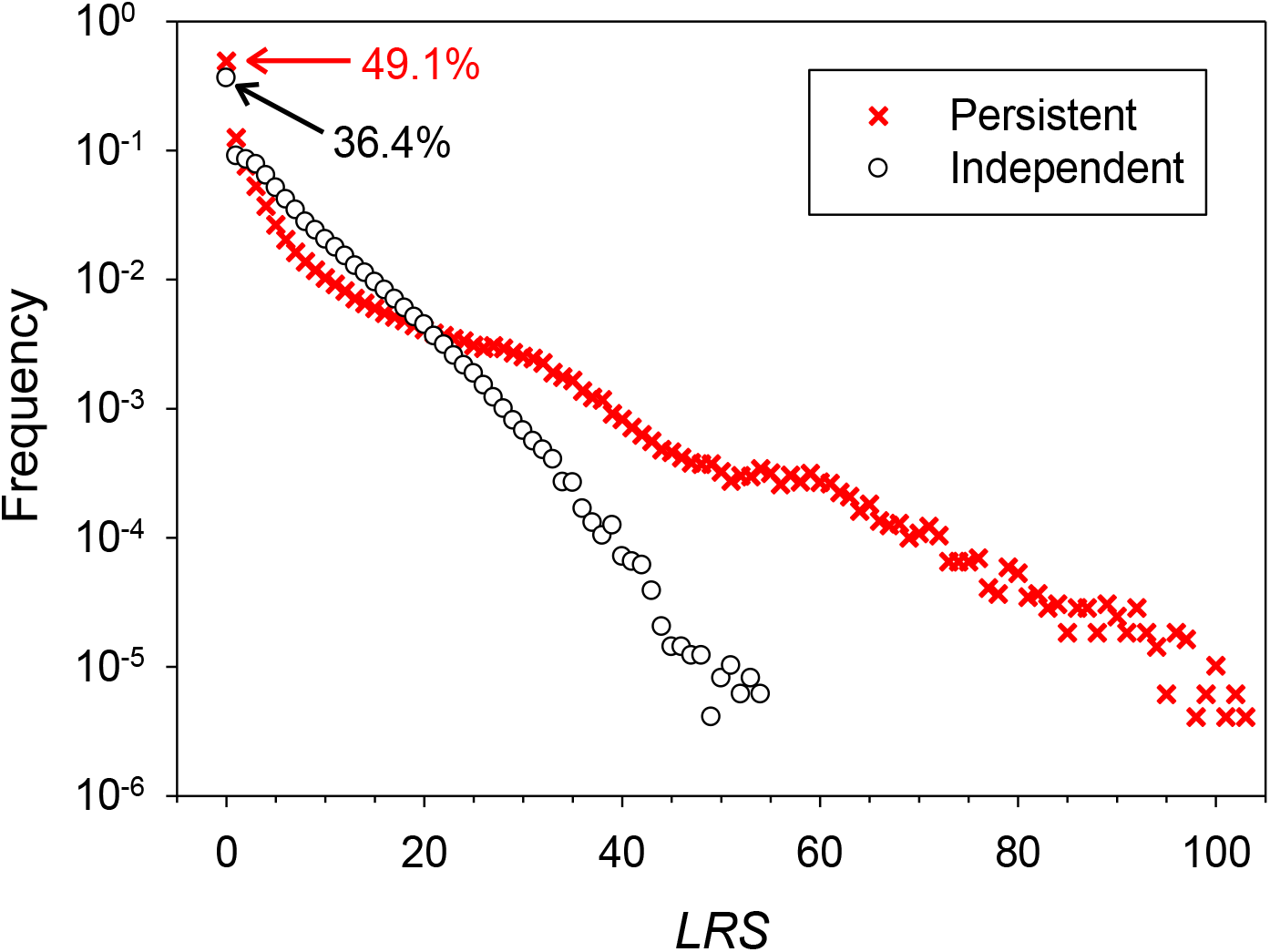
Distribution of lifetime reproductive success (*LRS*) for populations simulated using the vital rates in Table 2. Results are combined across 1000 replicates. In each replicate, *LRS* was computed for each of 490 males in a cohort that survived to age 3. Open circles show results for simulations in which parental weights were randomly shuffled each year; red Xs show results for simulations in which individuals retained their weights throughout their reproductive lifespan. Numbers with arrows (49.1%; 36.4%) indicate the percentages of individuals in a cohort that survived to age at maturity (3) but produced zero offspring across their lifetime, for persistent and independent simulations, respectively. Not included here are the ~51% of individuals in each cohort that died before reaching age 3 (see *N_x_* column in Table 1).

The examples in Tables 1 and 2 calculated means and variances of offspring number only for adults. If juveniles (all of which produce 0 offspring) had also been included, the overall means would have been lower and the variance-to-mean ratios would have been higher. Hence, to avoid confusion resulting from apples-and-oranges comparisons, it is important to clarify the groups of individuals that are included in analysis of reproductive success.

### 3.3 Simulating demography and genotypes

Because the novel features of *TheWeight* algorithm all involve population demography (specifically, how parents are chosen), the previous examples have all focused on that aspect of computer simulations. Nevertheless, most users are interested in (and the richest insights are gained from) joint modeling of demography and genetics, so code is also provided to illustrate one scenario that uses *TheWeight* to simulate reproduction in males and females and track multilocus genotypes over time (see Supporting Information for simulation details). Results show that (1) realized demographic parameters agreed well with expected values based on the specified vital rates (Tables S3 and S4), and that (2) the rate of decline of heterozygosity in the modeled population closely tracked the expected rate, based on expected *N_e_* calculated from the input data (Figure 4). Code for this example was also implemented in SLiM (Haller and Messer 2019), which produced comparable results (Figures S1 and S2).

**Figure 4.**
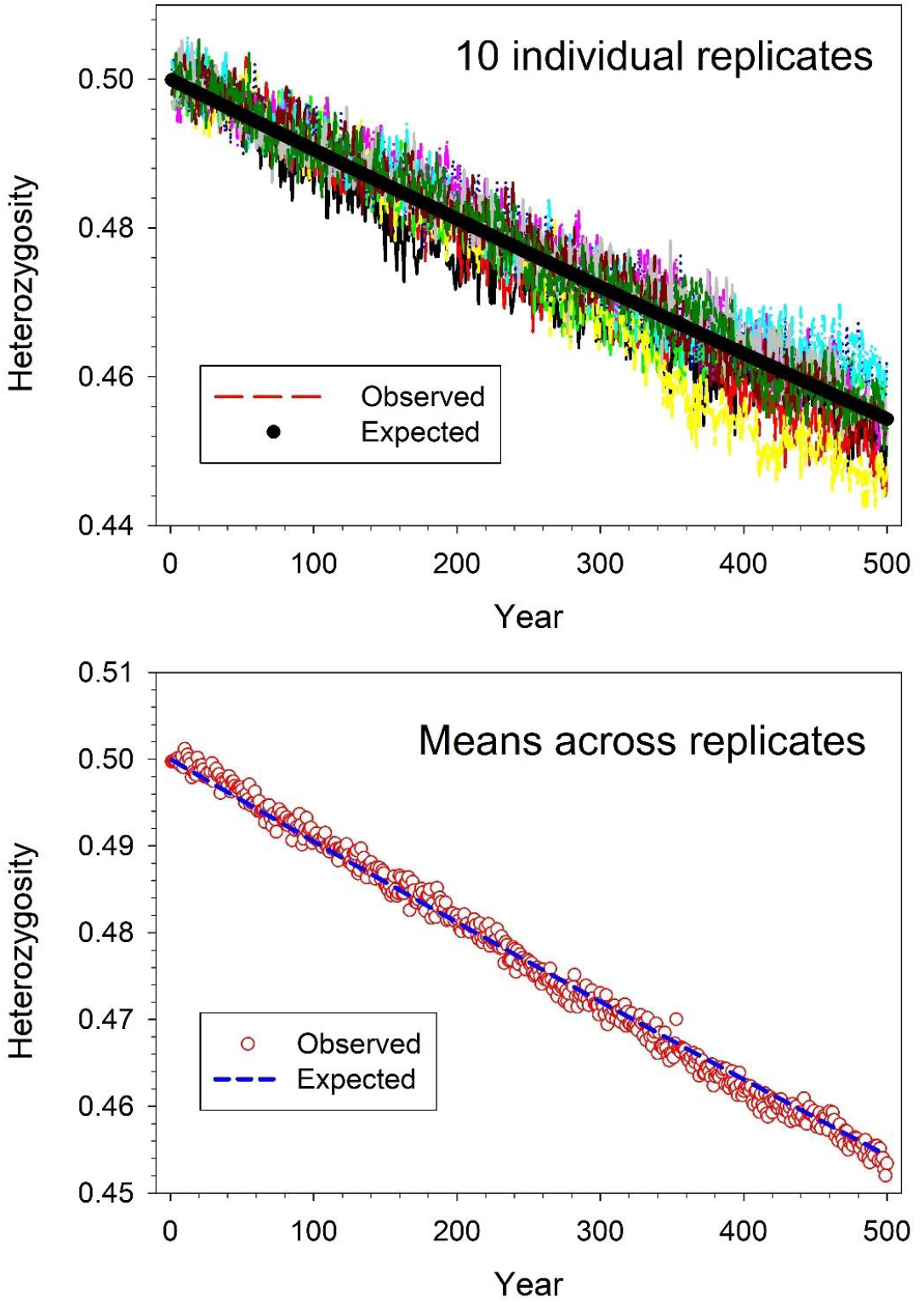
Observed and expected rate of loss of heterozygosity in age-structured populations modeled with *THEWEiGHT* algorithm. Vital rates for males are shown in Table 1 and those for females in Table S2. Generation length was 5.218 years, so each replicate of 500 years covered almost 96 generations. Each year, observed heterozygosity was averaged across 100 diallelic loci, and results are shown for each of 10 replicates (top) or averaged across replicates (bottom). The blue ‘expected’ line in the bottom panel was calculated using Equation S3 in Supporting Information, which provides more details regarding these simulations.

## 4 DISCUSSION

### 4.1 Recap of simple rules for implementing *TheWeight*

Age structure is one of the most common ways that most real populations depart from Wright-Fisher assumptions. *TheWeight* algorithm can be used to precisely control the distribution of reproductive success in age-structured populations by following a few simple rules.

1. For each age and sex, determine the desired degree of reproductive skew by specifying the value of 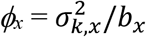, where *b_x_* is age-specific fecundity from a standard life table. Both *b_x_* and *ϕ_x_* should be scaled to their expected values in a stable population.
2. Calculate the magnitude of variation in parental weights that can be expected to produce the desired level of reproductive skew. The required level of variation in weights is 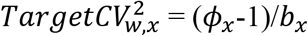.
3. Identify the parameter of the Poisson distribution that has the desired properties for the parental weights. The desired Poisson parameter, which is both the mean and the variance of the distribution, is 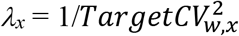.
4. Use a random Poisson process with parameter *λ_x_* to generate vectors of parental weights, until a vector is found with realized 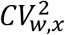 within an acceptable range of the target.
5. Repeat steps 1-4 for all ages and both sexes, if applicable.
6. If the one-step method is used for choosing parents, rescale the age-specific ***w_x_*** vectors so that their means equal the age-specific fecundities, *b_x_*. This is necessary to ensure that age-specific reproductive success matches the desired distribution, and that the realized generation length matches the target value. This rescaling can be accomplished by multiplying each vector ***w_x_*** by an appropriate scaling factor. Multiplying a vector by a constant changes the variance as well as the mean but does not change the coefficient of variation.
7. If persistent individual differences are modeled, allow individuals that survive each year to retain their weights rather than being assigned random new ones. If *b_x_* and/or *ϕ_x_* change with age, the vectors of weights for the survivors have to be adjusted to match the new age-specific 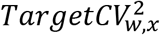. This can be accomplished by adding an age-specific constant *C_x_* to each weight, which has no effect on the variance but changes the mean and hence 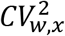. Supporting Information includes some detailed suggestions for implementing this option.

### 4.2 Model comparison

The Wright-Fisher model has two distinctive features: equiprobability and independence. Under WF, every individual is equally likely to be the parent of any given offspring, so offspring can be thought of as assigned to parents randomly and with equiprobability. Furthermore, each assignment is independent, in the sense that the outcome does not depend on results of any previous assignments. In R (for example), the WF process can be modeled using the ‘sample’ function, with *size* = the number of offspring and sampling done from a vector of parental names or IDs. Under the defaults, sampling is equiprobable and done with replacement and hence independent.

In the generalized WF model (Waples 2020), the assignments are still independent (within time steps at least) but parental assignment probabilities are allowed to vary according to the vector of parental weights, ***w***. This only requires adding a ‘prob = w’ argument to the ‘sample’ function. Unequal parental weights lead to overdispersed variance in offspring number compared to the Poisson expectation.

Another common approach to modeling overdispersed variance is to randomly assign numbers of offspring to parents based on a negative binomial or gamma distribution that can be parameterized to generate a target *μ_k_* and 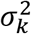 (e.g., Anderson and Dunham 2005). This approach is fast and simple but has some limitations compared to *TheWeight*. First, when offspring numbers are chosen randomly from a distribution, it is not easy (as it is in *TheWeight*) to relate expected reproductive success to traits or attributes of individuals. Second, this same limitation makes it difficult to control the degree of correlation in expected individual reproductive success over time, which can easily be accomplished using *TheWeight*.

### 4.3 Modeling selection

The standard WF model assumes a very large gamete pool equally produced by *N* parents and random survival of zygotes until the next generation of *N* offspring is produced, leading to *μ_k_* = 2. The generalized WF model also assumes random survival of zygotes but allows for unequal parental contributions to the initial gene pool and allows mean offspring number to vary; this latter feature is important for modeling seasonal reproduction in iteroparous species. As a consequence, *TheWeight* algorithm is well suited for modeling fertility selection but does not explicitly model viability selection. However, the latter can also easily be incorporated by inserting an episode of selective mortality that relates probability of survival to a trait of interest.

### 4.4 Null models

Null models are increasingly important in evolutionary biology, including for the analysis of reproductive success (van Daalen and Caswell 2017; Tuljapurkar et al. 2020). A variety of null models are easy to implement using *TheWeight*. For seasonal reproduction, one obvious null model is to treat all adults as a single WF population, which can be accomplished by setting fecundity to be constant with age and all ϕ = 1. For *LRS*, randomly scrambling parental weights each year replicates a null model of independence of reproductive success over time. Comparison with results for this null model can dramatically illustrate the consequences of a lack of independence (see Figure 3).

## Acknowledgments

I thank Eric Anderson, Tom Reed, Ryan Waples, and two anonymous reviewers for useful comments and suggestions. Ryan K. Waples implemented the model in SLiM. The author declares no conflict of interest.

## Data availability statement

This study did not generate any new empirical data, except by simulation. Code to conduct the simulations and analyses described here is available on Zenodo at https://doi.org/10.5281/zenodo.6585639.

