## Supplemental info for "*TheWeight*: A simple and flexible algorithm for simulating non-ideal, age-structured populations"

Computer code to execute the examples described here and in the text is available on Zenodo at https://doi.org/10.5281/zenodo.6585639.

**SI I. Exact solution for binomial variance in Equation 1**.

From Equation 1 (main text), the discrete-generation equation for inbreeding $N_e$ (which also applies to the inbreeding effective number of breeders per year, $N_b$) is

$$\text{Inbreeding } N_e = \frac{\mu_k N-1}{\mu_k-1+\phi}, \tag{S1}$$

where $\phi = \sigma_k^2/\mu_k$ is the ratio of variance to mean offspring number. In the main text, it was pointed out that if one ignores the "-1" in the numerator and uses the Poisson approximation that $\phi$ = 1under random reproductive success, the above equation reduces to

$$\text{Inbreeding } N_e \approx \frac{\mu_k N}{\mu_k} = N.$$

In fact, this relationship holds more generally, as can be shown using the exact (binomial) variance in offspring number ($\sigma_k^2 = \frac{\mu_k(N-1)}{N}$) that applies to random, Wright-Fisher reproduction, in which case $\phi = \sigma_k^2/\mu_k = (N-1)/N$:

$$\text{Inbreeding } N_e = \frac{\mu_k N-1}{\mu_k-1+\phi} = \frac{\mu_k N-1}{\mu_k-1+(N-1)/N} = \frac{\mu_k N-1}{\mu_k-1/N} = \frac{\mu_k N-1}{(\mu_k N-1)/N} = N. \tag{S2}$$

Equation S2 holds generally for inbreeding $N_e$ but is true for variance effective size only when population size is constant (hence $\mu_k$ = 2).

**SI II. Simulating demography and multilocus genotypes.**

Results shown in Figure 4 were generated from simulations that modeled both demography and genetics in both sexes. Vital rates for males used the $s_x$, $b_x$, and $\phi_x$ values shown in Table 1 (so both $b_x$, and $\phi_x$ increased with age in males). Female survival was the same as for males (constant at 0.7/year), but otherwise female vital rates were simpler, fecundity being constant with age and reproductive success random among individuals of the same age, so all $\phi_x$ = 1 (Table S2). In this modeled scenario, there were no persistent individual differences in reproductive success across years, so surviving individuals received new random weights each year. Age at maturity was 3 in both sexes, so all juveniles received 0s for their weight. For males, the one-step approach was used for implementing *THEWEIGHT* algorithm. Within each age class, weights were first randomly generated to meet the age-specific Target $CV_{w,x}^2$, and then each weight was multiplied by an age-specific scaling factor (as shown in Table 1) to ensure that $\bar{w}_x$ = $b_x$ for each age. Modeling female reproduction was simpler: because $b_x$ was fixed and reproductive success was random within ages, each year every adult female had an equal opportunity to produce offspring. As a consequence, all adult females had identical weights that did not change with age (in this example all $\boldsymbol{w}$ = 1, but any constant positive value would produce the same result).

The model assumed a fixed cohort size of 400 age-1 offspring, with each offspring having an equal probability of being male or female. With an average of $N_1$ = 200 yearlings of each sex in each cohort, the expected numbers alive at subsequent ages are given by $E(N_x) = N_1 l_x$, as shown in Table S2. In each replicate, the population was initiated with these expected numbers of males and females of each sex. Subsequently, random variation in sex ratio and actual survival rates created variation in realized $N_x$ values independently in both sexes.

In addition to its sex, unique ID, age, and parental weight, associated with each individual were genotypes at $L$ = 100 diallelic loci. At each locus, these genotypes were coded 0/1/2 to indicate the number of copies of the focal allele. Thus, genotypes "0" and "2" represent alternative homozygotes (0 or 2 copies of the focal allele), whereas genotype "1" indicates a heterozygote. For simplicity in tracking the rate of loss of heterozygosity, in the initial population all individuals were coded "1" at every locus, so all loci started with equal allele frequencies $P$ = 1-$P$ = 0.5. Following an episode of random mating, offspring genotypes were in expected Hardy Weinberg proportions, with heterozygosity varying randomly around 0.5. Mutation was not modeled, so the expected rate of loss of heterozygosity over time was calculated from the relationship (Crow and Kimura 1970)

$$H_t = H_0[1 - 1/(2N_e)]^t,$$

where $H_0$ and $H_t$ are heterozygosities at times 0 and $t$, respectively, and $t$ is elapsed time in generations. For the vital rates used, generation length (average age of parents) was 5.218 years (4.84 for females and 5.59 for males) and generational $N_e$ was 500.3, using the *AgeNe* method of Waples et al. (2011). In the simulations, therefore, the expected heterozygosity at year $Y$ was calculated as

$$H_Y = 0.5[1 - 1/1000.6]^t \ , \qquad \text{(S3)}$$

where $t = Y/5.218$ is elapsed number of generations. Each year in each replicate, observed heterozygosity was calculated as the fraction of "1" genotypes in the new cohort. To generate the results shown in Figure 4, 10 separate replicates were run for 500 years, and each year in each replicate mean heterozygosity was averaged across 100 loci. Although random differences in the multigeneration pedigree created substantial variation in mean heterozygosity across replicates (top panel of Figure 4), heterozygosity averaged across replicates agreed very closely with the theoretical expectation (bottom panel of Figure 4). Code to conduct these simulations, including an example implementation in SLiM, is available on Zenodo.

The expected $N_e$ = 500.3 from *AgeNe* assumes that population demography each generation exactly follows the parametric vital rates in the input file(s). The simulation code records the pedigree of offspring produced each year, which facilitates demographic analysis of the data. As shown in Table S3, harmonic mean $N_e$ recorded in the simulations (496.3) was very close to the target value, as were other indices of seasonal and lifetime reproductive success. The same was true for a comparison of observed and expected age-specific $\phi$ values (Table S4), although in the oldest age class (10) the deviations from expected values were slightly higher.

This reflects the fact that, with only eight 10-year-olds of each sex expected to be alive each year (from the $N_x$ column of Table S2), random variation in sex ratio and survival caused large proportional changes over time in the realized $N_{10}$ values. R code to analyze the pedigree and produce summaries similar to those shown in Tables S3 and S4 is also available on Zenodo.

### SI III. Notes on generating weights

The methods for generating weights described in the text and implemented in the examples are generally effective in producing vectors of weights with the desired properties. Below are some suggestions for fine tuning of the weights that can be useful in specific situations.

#### SI III.A. Small numbers of individuals

Raw weights generated by a random Poisson process are all integers. When the number of potential parents is relatively large, the distribution of possible $CV_w^2$ values is almost continuous, which means that Monte Carlo methods can generally find a vector of weights with a $CV_w^2$ very close to the target value. For example, a tolerance of 1% was used for accepting weight vectors in the examples shown in Tables 1 and 2. As the number of potential parents declines, the distribution of potential $CV_w^2$ values becomes more coarse grained, and it might not always be possible to achieve a vector whose $CV_w^2$ lies within the specified tolerance. This scenario is most likely to occur for older age classes in small populations of long-lived species, as occurred in the example shown in Figure 4. This problem is exacerbated if age-specific $\phi$ is high, in which case target $CV_w^2$ will also be large. This generally requires that most parents receive a weight of zero and only a small percentage get positive weights, which in turn constrains the range of possible solutions. To deal with this issue, one can use a sliding scale for the tolerance criterion: very demanding (one or a few percent) for $N_x$ values above a certain threshold, with more relaxed tolerance as $N_x$ declines. The GetWeights function for the demography+genetics simulations used a sliding scale like this: tolerance was 2% for $N_x > 20$, 5% for $20 \geq N_x > 10$, and $(100/N_x)$% for $N_x < 10$. Relaxing the tolerance allows greater variation among replicates in the vector of weights but has relatively little effect on mean demographic parameters across many replicates. The user can experiment with the tolerance criterion to achieve desired results.

The most extreme parental-weighting scenario occurs when only one individual has a positive weight and all others have zero. In this situation, $CV_w^2$ is the same regardless what the weight of the super parent is, and the maximum possible $CV_w^2$ is constrained by $N_x$. To see this, let one parent have weight $X$ and $n$-1 parents have weight 0. Then the mean weight is $\bar{w} = X/n$, the variance of the weights is

$$\sigma_w^2 = \frac{X^2}{n} - \left(\frac{X}{n}\right)^2 = \frac{X^2(n-1)}{n^2} = (X/n)^2(n-1) \qquad ,$$

and the squared coefficient of variation of the weights is

$$CV_w^2 = \sigma_w^2/\bar{w}^2 = \frac{(X/n)^2(n-1)}{(X/n)^2} = n - 1 \qquad .$$

Because $CV_w^2$ can never be larger than $N_x$ -1, it is important to include a condition in the GetWeights function that identifies this scenario, bypasses the *rpois* routine, and returns a vector with one positive weight and $N_x$ -1 zero weights. This feature is incorporated in the R code for the demography+genetics example.

**SI III.B. Persistent individual differences**

Figure 3 illustrates the profound effect that persistent individual differences can have on the distribution of lifetime reproductive success. As discussed in the main text, persistent differences can be modeled by having individuals retain their weights across their adult lifespans. As illustrated in Table 2, if the target $CV_w^2$ varies with age, existing weights of survivors can be adjusted by adding an appropriate age-specific constant *C* to all individuals of a given age.

A potential problem occurs when the value of *C* is <0; since all weights have to be non-negative, adding a negative constant to the weights of all individuals within an age class can produce some weights with impermissible values. Negative values of *C* occur when $Q$ > 1, where $Q$ is the ratio of the Target $CV_w^2$ in the current year to target $CV_w^2$ in a previous year. For age $x$, Taget $CV_{w,x}^2$ = ($\phi_x$-1)/$b_x$, so $Q$ > 1 can occur when $\phi$ increases with age more rapidly than does fecundity. A simple solution to this potential problem is to convert all negative weights to 0. However, this will reduce the variance of the weights and hence affect the realized $CV_w^2$. If the effect is substantial enough to distort results from the desired patterns of age-specific reproductive success, the user can implement a simple 'spreading' routine to increase the variance and $CV_w^2$ of the weights:

(1) Define a **Multiplier** vector with a range of positive values greater than and less than 1. Here is one example: **Multiplier** = c(0.1, 0.2, 0.5, 1, 2, 5, 10).
(2) Multiply the weight of each individual by a randomly-chosen value from **Multiplier**.
(3) Step 2 will increase the standard deviation of the weights more than the mean (or reduce the mean more than the standard deviation) and hence increase $CV_w^2$. The age-specific mean weight can then be adjusted by multiplying all weights by an appropriate constant.

The user can tune this 'spreading' function to produce the desired magnitude of effect.

Cited here but not in main text:

Crow, JF, and M Kimura. 1970. An introduction to population genetics theory. Harper and Row, New York.

Table S1. As in Table 1 (main text), but with all adult $\phi_x$ set to 1.0 to model random reproductive success of individuals of the same age and sex. Fecundity still increases with age. The bottom part of the table compares results (means across 1000 replicates) for all adults reproducing each year, for the scenario in Table 1 and this scenario with $\phi_x$ =1.

| | | | | Observed | | |
|---|---|---|---|---|---|---|
| **Age** | $N_x$ | $b_x$ | $\phi_x$ | $\phi_x$ | $\bar{w}_x$ | $CV^2_{w,x}$ |
| 1 | 1000 | 0 | | | | |
| 2 | 700 | 0 | | | | |
| 3 | 490 | 0.805 | 1.00 | 1.001 | 0.805 | 0.000 |
| 4 | 343 | 1.073 | 1.00 | 1.001 | 1.073 | 0.000 |
| 5 | 240 | 1.342 | 1.00 | 0.998 | 1.342 | 0.000 |
| 6 | 168 | 1.610 | 1.00 | 0.999 | 1.610 | 0.000 |
| 7 | 118 | 1.878 | 1.00 | 1.002 | 1.878 | 0.000 |
| 8 | 82 | 2.146 | 1.00 | 0.999 | 2.146 | 0.000 |
| 9 | 58 | 2.415 | 1.00 | 1.004 | 2.415 | 0.000 |
| 10 | 40 | 2.683 | 1.00 | 0.993 | 2.683 | 0.000 |

| | **Table 1** | **Table S1** |
|---|---|---|
| $\mu_k$ | 1.30 | 1.30 |
| $\sigma^2_k$ | 5.00 | 1.56 |
| $\phi$ | 3.85 | 1.20 |
| $\bar{w}$ | 1.30 | 1.30 |
| $\sigma^2_w$ | 3.70 | 0.26 |
| $CV^2_w$ | 2.19 | 0.15 |

Table S2. Female vital rates for the scenario depicted in Figure 4, which jointly modeled demography and tracked multilocus genotypes. Male vital rates for this scenario are shown in Table 1. $s_x$ is probability of surviving from age $x$ to $x+1$; $N_x$ is the expected number alive at age $x$; $b_x$ is mean fecundity at age $x$, scaled to constant population size; $B_x$ is the expected number of offspring produced each year by individuals of age $x$; $\phi_x = \sigma^2_{k,x}/b_x$ is the ratio of variance to mean offspring number for individuals of age $x$. In this scenario, a fixed total of 400 age-1 offspring are produced each year, each with equal probability of being male or female, so $E(N_1)$ = 200 for both sexes. In the simulations, stochasticity associated with sex determination and survival caused $N_x$ to vary randomly in both sexes.

| **Age** | $s_x$ | $N_x$ | $b_x$ | $B_x$ | $\phi_x$ |
|---|---|---|---|---|---|
| 1 | 0.7 | 200 | 0 | 0 | NA |
| 2 | 0.7 | 140 | 0 | 0 | NA |
| 3 | 0.7 | 98 | 1.299 | 127 | 1 |
| 4 | 0.7 | 69 | 1.299 | 89 | 1 |
| 5 | 0.7 | 48 | 1.299 | 62 | 1 |
| 6 | 0.7 | 34 | 1.299 | 44 | 1 |
| 7 | 0.7 | 24 | 1.299 | 31 | 1 |
| 8 | 0.7 | 16 | 1.299 | 21 | 1 |
| 9 | 0.7 | 12 | 1.299 | 15 | 1 |
| 10 | 0 | 8 | 1.299 | 10 | 1 |
| | | | | 400 | |

Table S3. Summary of annual and lifetime demographic data for the demography+genetics scenario, whose results are shown in Figure 4. Observed values are means (Generation length, Lifetime variance in offspring number, $V_{k*}$) or harmonic means ($N_b$, $N_e$) across 500 years in one replicate. Expected values were calculated from AGENE (Waples et al. 2011) using the input data shown in Tables 1 (for males) and S2 (for females). The effective number of breeders ($N_b$) was calculated for years 21-480, after excluding the first and last 20 years (= 2 maximum lifespans) of data. Lifetime measures were calculated for cohorts born in years 21-480.

| | | Female | Male | Overall |
|---|---|---|---|---|
| **Observed** | $N_b$ | 307.1 | 95.4 | 291.2 |
| | Generation | 4.83 | 5.56 | 5.19 |
| | Lifetime $V_{k*}$ | 10.16 | 19.33 | 14.75 |
| | $N_e$ | 317.2 | 207.3 | 496.3 |
| **Expected** | $N_b$ | 313.4 | 96.2 | 294.4 |
| | Generation | 4.84 | 5.59 | 5.22 |
| | Lifetime $V_{k*}$ | 10.13 | 19.25 | 14.69 |
| | $N_e$ | 319.4 | 210.6 | 500.3 |

Table S4. As in Table S3, but summarizing age- and sex-specific values of $\phi$ for seasonal reproduction in adults. Expected $\phi_x$ for males is from Table 1. For females, the fixed input value of $\phi = 1$ in Table S2 indicates random reproductive success among adults, in which case the expected value of $\phi_x$ is actually $(N_x\text{-}1)/N_x$, and these expected values (based on the $N_x$ vector in Table S2) are shown below. Observed values of $\phi_x$ are means across years 21-480 in one replicate simulation. $\phi$ was only calculated for replicates in which the number of surviving individuals of the respective age was >5, which resulted in reduced replication for the oldest age classes (especially age10, for which the expected number of survivors of each sex was only 8).

| | Male | | Female | |
|---|---|---|---|---|
| Age | Expected | Observed | Expected | Observed |
| 3 | 3.00 | 3.03 | 0.990 | 0.998 |
| 4 | 3.25 | 3.27 | 0.986 | 0.988 |
| 5 | 3.50 | 3.51 | 0.979 | 0.966 |
| 6 | 3.75 | 3.78 | 0.971 | 0.952 |
| 7 | 4.00 | 3.95 | 0.958 | 0.947 |
| 8 | 4.25 | 4.16 | 0.938 | 0.924 |
| 9 | 4.50 | 4.71 | 0.917 | 0.907 |
| 10 | 4.75 | 5.02 | 0.875 | 0.978 |

Table S5. Vital rates for alternative life tables for 5- and 20-year lifespans. These vital rates were used in the simulations whose results are shown in Figure S2. In both scenarios, total cohort size of age-1 offspring was $2*N_1 = 400$, where $N_1$ is expected number of yearling offspring of each sex. In the simulations, the actual sex ratio varied randomly around the 1:1 expectation (see Section SI II for details). Age specific $b_x$ and $\phi_x$ are scaled to values that will produce a stable population.

| Age | FemaleSx | Femalebx | Femalephi | MaleSx | Malebx | Malephi |
|---|---|---|---|---|---|---|
| 1 | 0.5 | 0.561 | 5 | 0.5 | 0.561 | 5 |
| 2 | 0.5 | 1.123 | 5 | 0.5 | 1.123 | 5 |
| 3 | 0.5 | 1.684 | 5 | 0.5 | 1.684 | 5 |
| 4 | 0.5 | 2.246 | 5 | 0.5 | 2.246 | 5 |
| 5 | 0 | 2.807 | 5 | 0 | 2.807 | 5 |

| Age | FemaleSx | Femalebx | Femalephi | MaleSx | Malebx | Malephi |
|---|---|---|---|---|---|---|
| 1 | 0.85 | 0 | 2 | 0.85 | 0 | 2 |
| 2 | 0.85 | 0 | 2 | 0.85 | 0 | 2 |
| 3 | 0.85 | 0 | 2 | 0.85 | 0 | 2 |
| 4 | 0.85 | 0 | 2 | 0.85 | 0 | 2 |
| 5 | 0.85 | 0.621 | 2 | 0.85 | 0.621 | 2 |
| 6 | 0.85 | 0.621 | 2 | 0.85 | 0.621 | 2 |
| 7 | 0.85 | 0.621 | 2 | 0.85 | 0.621 | 2 |
| 8 | 0.85 | 0.621 | 2 | 0.85 | 0.621 | 2 |
| 9 | 0.85 | 0.621 | 2 | 0.85 | 0.621 | 2 |
| 10 | 0.85 | 0.621 | 2 | 0.85 | 0.621 | 2 |
| 11 | 0.85 | 0.621 | 2 | 0.85 | 0.621 | 2 |
| 12 | 0.85 | 0.621 | 2 | 0.85 | 0.621 | 2 |
| 13 | 0.85 | 0.621 | 2 | 0.85 | 0.621 | 2 |
| 14 | 0.85 | 0.621 | 2 | 0.85 | 0.621 | 2 |
| 15 | 0.85 | 0.621 | 2 | 0.85 | 0.621 | 2 |
| 16 | 0.85 | 0.621 | 2 | 0.85 | 0.621 | 2 |
| 17 | 0.85 | 0.621 | 2 | 0.85 | 0.621 | 2 |
| 18 | 0.85 | 0.621 | 2 | 0.85 | 0.621 | 2 |
| 19 | 0.85 | 0.621 | 2 | 0.85 | 0.621 | 2 |
| 20 | 0 | 0.621 | 2 | 0 | 0.621 | 2 |

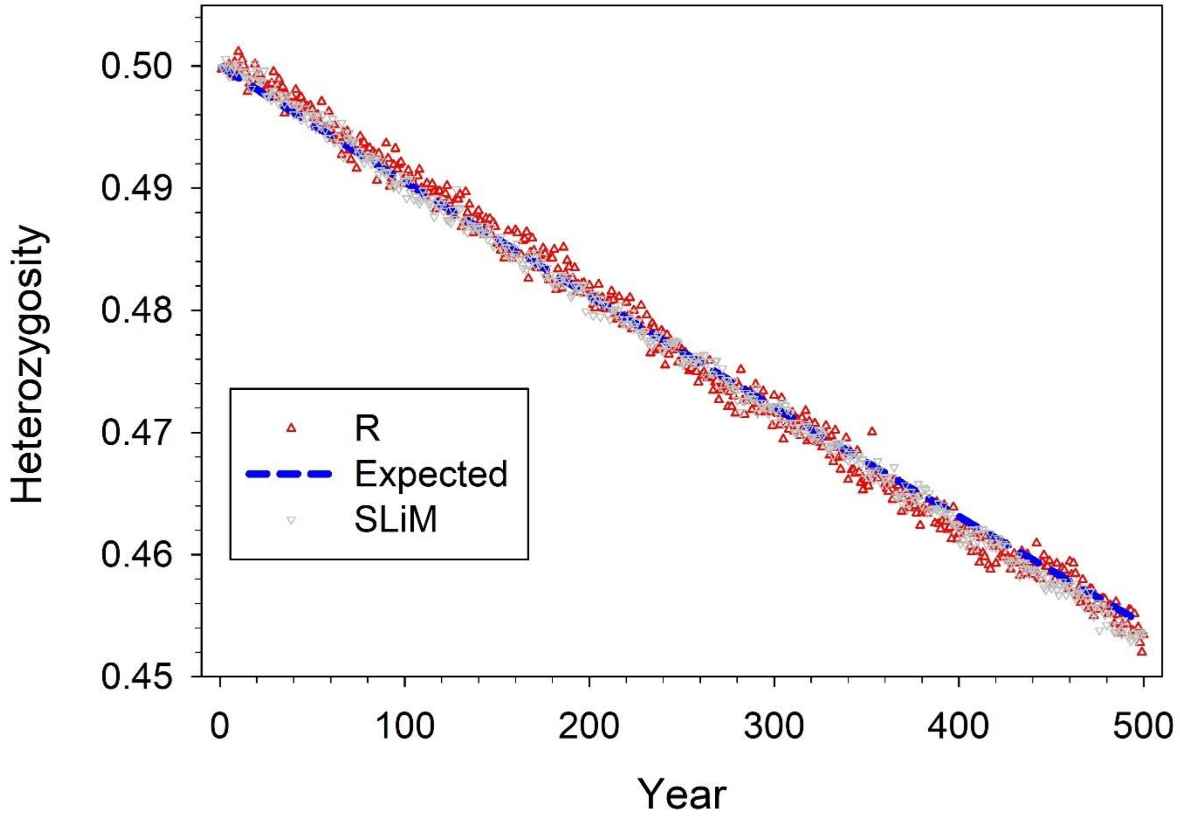


Figure S1. As in Figure 4, except adding results for the implementation in SLiM. These simulations tracked loss of heterozygosity over time in a population with vital rates for males as shown in shown in Table 1 and vital rates for females as shown in Table S2. Each year, observed heterozygosity was averaged across 100 diallelic loci, and then averaged across 10 different replicates. The blue 'expected' line was calculated for these vital rates using Equation S3 in Supporting Information.

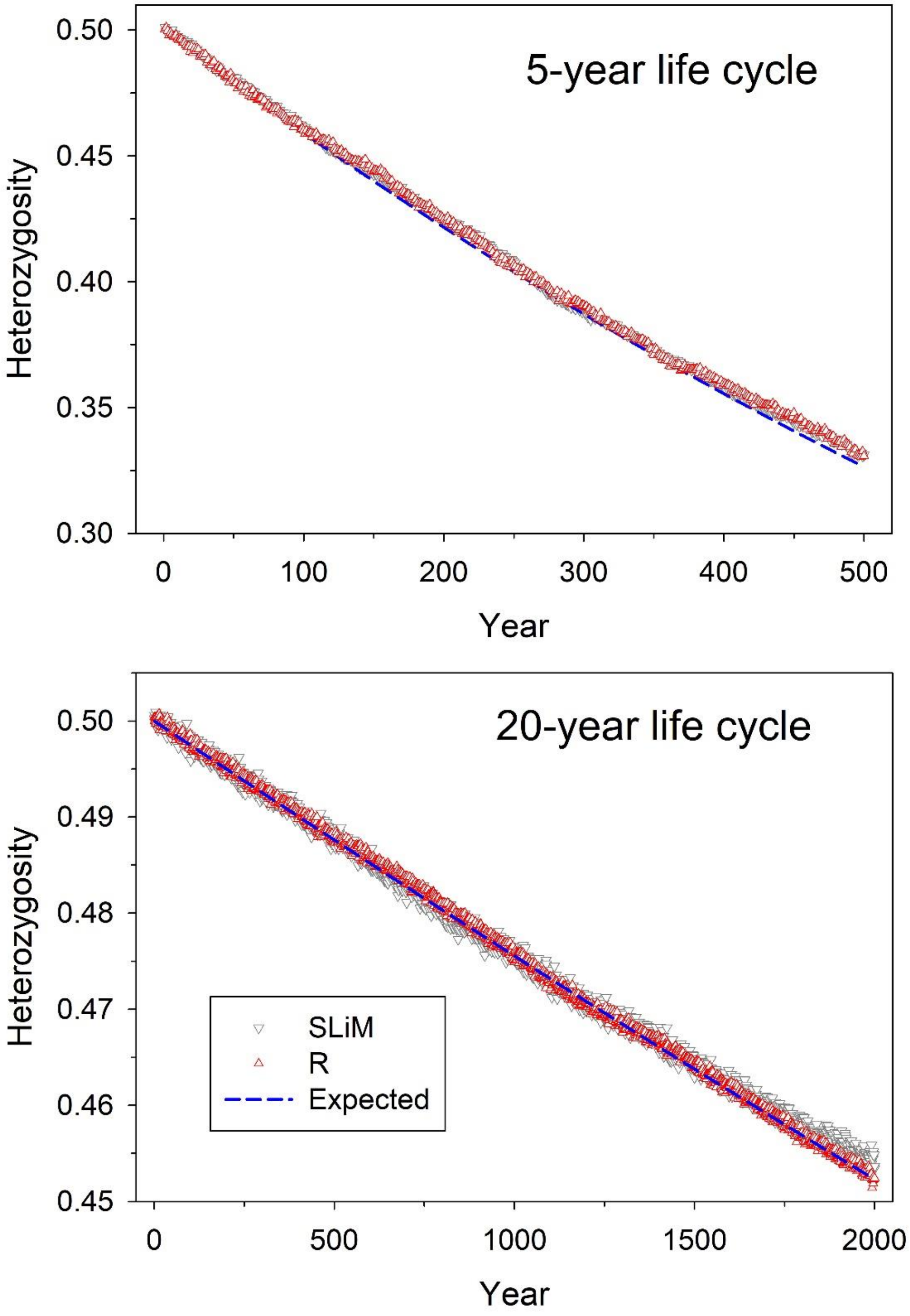


Figure S2. As in Figure S1, but for simulations using the alternative life tables for 5- and 20-year lifespans shown in Table S5.
